## Supplemental data for "Hydrogen sulfide (H_2_S) coordinates redox balance, carbon metabolism, and mitochondrial bioenergetics to suppress SARS-CoV-2 infection"

**Supplementary figure:1**

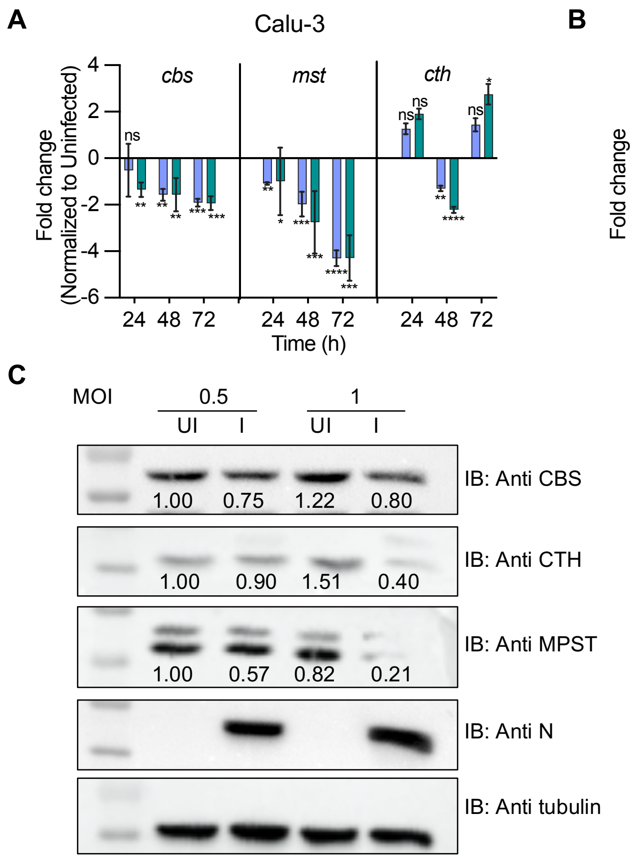


**Supplementary Figure 1: (A)** Time-dependent changes in expression of *cbs*, *mst* and *cth* during SARS-CoV-2 (HK variant) replication in Calu-3 cells by RT-qPCR. **(B)** Time-dependent changes in expression of *cbs*, *mst* and *cth* during SARS-CoV-2-HK replication in HEK-ACE2 cells by RT-qPCR. **(C)** Protein levels of CBS, CTH and MPST during SARS-CoV-2-HK replication in HEK-ACE2 cells.

**Supplementary figure:2**

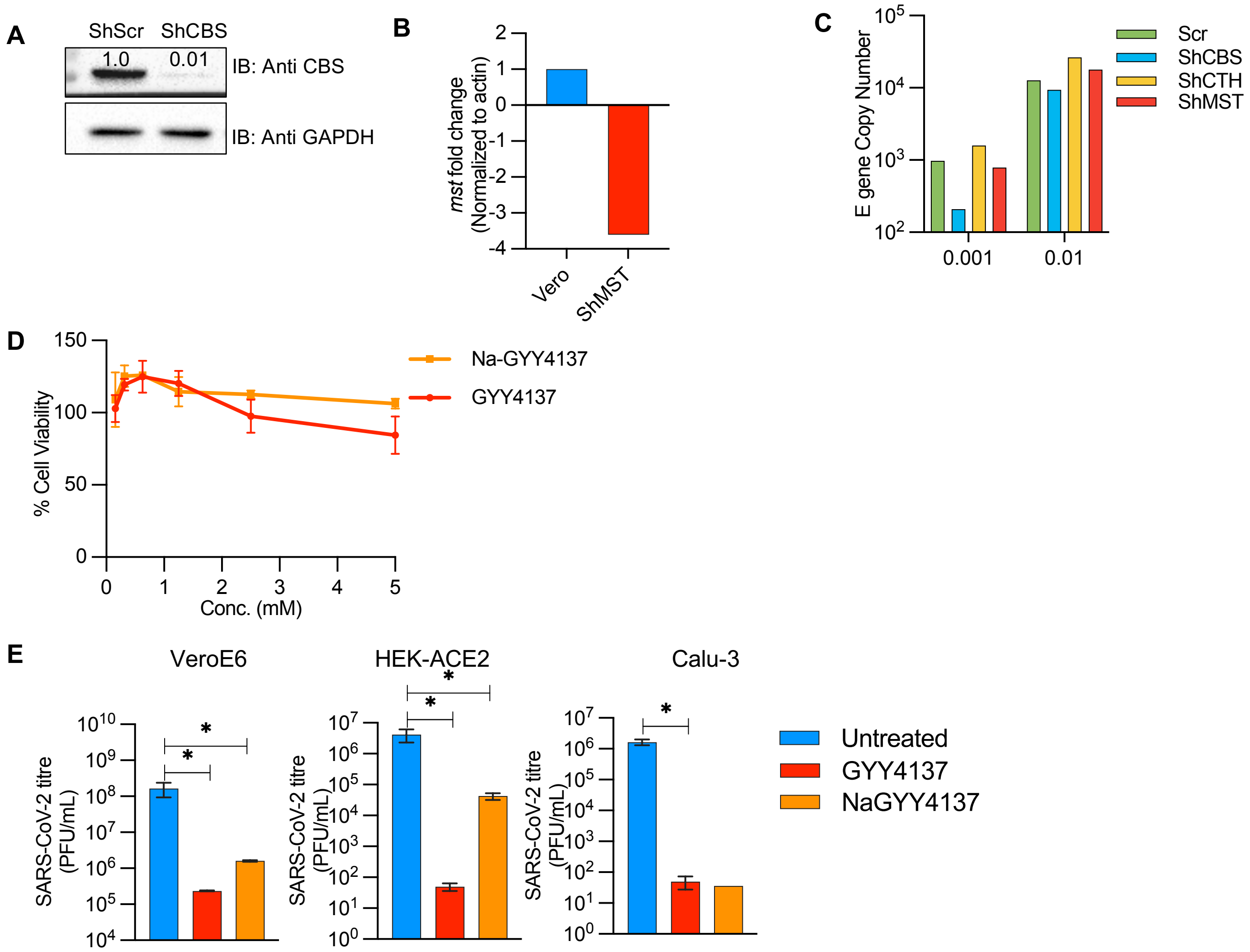


**Supplementary Figure 2: (A)** Knockdown confirmation of CBS in VeroE6 cells by western blotting. **(B)** Knockdown confirmation of MST in VeroE6 cells by RT-qPCR. **(C)** SARS-CoV-2 viral load in knockdown VeroE6 cells (ShCBS/CTH/MST). **(D)** Viability of VeroE6 cells in the presence of GYY4137 and Na-GYY4137 at 48 h post treatment by MTT assay. **(E)** Plaque assay from culture supernatant of different treatment groups.

**Supplementary figure:3**

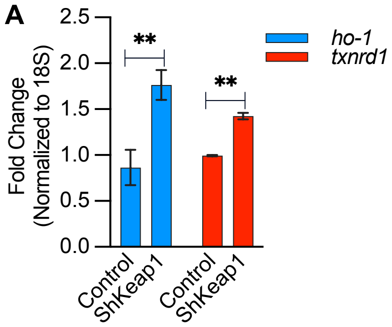


**Supplementary Figure 3: (A)** RT-qPCR analysis of Nrf2 genes in Vero-shKeap1 cells.

**Supplementary figure:4**

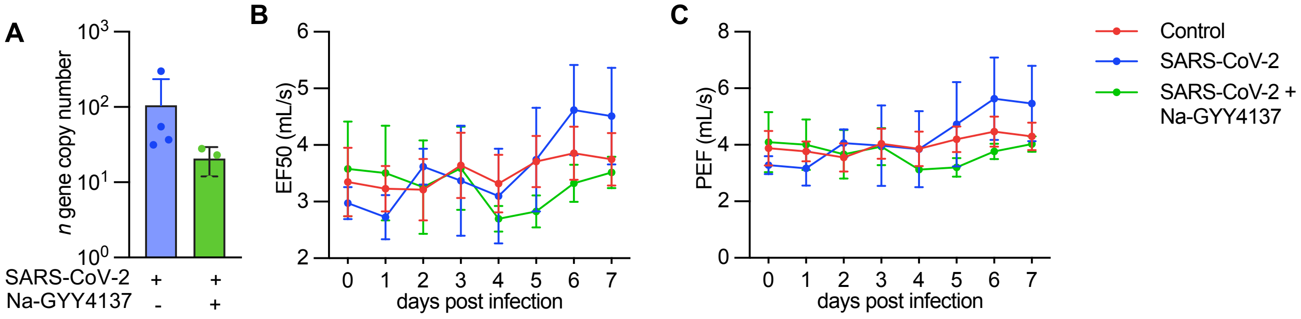


**Supplementary Figure 4: (A)** Viral load in SARS-CoV-2 infected mice in presence or absence of Na-GYY4137 measured by RT-qPCR at 7 day p.i. (**B**) Lung function parameters measured by whole body plethysmography.

**Supplementary table 1**: List of shRNA constructs used in the study

| Gene name | Accession number | Target Sequence | Code |
|---|---|---|---|
| *cbs* | NM_000071.2 | GCCGTCAGACCAAGTTGGCAAAGTC | C10 |
| *cth* | NM_001902.4, NM_153742.3 | GGCACCTCATTATCTTTCATAACT | E8 |
| *mst* | NM_021126 | AGAAGAAAGTGGACCTGTCTA | A8 |
| *keap1* | NM_012289.3, NM_203500.1 | CGGGAGTACATCTACATGCAT | A10 |

**Supplementary table 2**: List of primers used in the study

| Species | Gene | Primer | Sequence (5'-3') |
|---|---|---|---|
| Human | *actin* | Forward | ATGTGGCCGAGGACTTTGATT |
| | | Reverse | AGTGGGGTGGCTTTTAGGATG |
| | *cbs* | Forward | GCCTGAAGTGTGAGCTCTTG |
| | | Reverse | CACGATGATGCAGCGATAGC |
| | *cth* | Forward | TATTTACTCTGGCCGAGAGC |
| | | Reverse | TCCTCTAAGCCCACAGAAAG |
| | *mst* | Forward | GCCGCTTTCTTCGACATC |
| | | Reverse | TGGCGTCGTAGATCACG |
| | *b2m* | Forward | GCCCAAGATAGTTAAGTGGGATCG |
| | | Reverse | TCATCCAATCCAAATGCGGC |
| | *18s* | Forward | GGCCCTGTAATTGGAATGAGTC |
| | | Reverse | CGCTCCCAAGATCCAACTAC |
| | *gclc* | Forward | GGCACAAGGACGTTCTCAAGT |
| | | Reverse | CAAGGGTAGGATGGTTTGGG |
| | *ho-1* | Forward | TAGAAGAGGCCAAGACTGCG |
| | | Reverse | GGGCAGAATCTTGCACTTTGTT |
| | *txn* | Forward | CCCTCTCTGAAAAGTATTCCAACG |
| | | Reverse | TGGCTCCAGAAAATTCACCCA |
| | *txnrd-1* | Forward | ATGGGCAATTTATTGGTCCTCAC |
| | | Reverse | CCCAAGTAACGTGGTCTTTCAC |
| | *cat* | Forward | TTAATCCATTCGATCTCACC |
| | | Reverse | GGCGGTGAGTGTCAGGATAG |
| | *gsr* | Forward | CTTGCGTGAATGTTGGATGT |
| | | Reverse | GACCTCTATTGTGGGCTTGG |
| | *gpx-4* | Forward | GCCTTCCCGTGTAACCAGT |
| | | Reverse | GCGAACTCTTTGATCTCTTCGT |
| | *gpx-1* | Forward | CAACCAGTTTGGGCATCAG |
| | | Reverse | GTTCACCTCGCACTTCTCG |
| Mouse | *gpx-1* | Forward | GGTTCGAGCCCAATTTTACA |
| | | Reverse | CCCACCAGGAACTTCTCAAA |
| | *gpx-4* | Forward | ACGTCAGTTTTGCCTCATTG |
| | | Reverse | CTCCATGCACGAATTCTCAG |
| | *cat* | Forward | GGACGCTCAGCTTTTCATTC |
| | | Reverse | TTGTCCAGAAGAGCCTGGAT |
| | *gclc* | Forward | ACACCTGGATGATGCCAACGAG |
| | | Reverse | CCTCCATTGGTCGGAACTCTAC |
| | *txnrd-1* | Forward | AGTCACATCGGCTCGCTGAACT |
| | | Reverse | GATGAGGAACCGCTCTGCTGAA |
| | *tnfa* | Forward | CTGCACTTTGGAGTGATCGG |
| | | Reverse | TCAGCTTGAGGGTTTGCTAC |
| | *il-6* | Forward | TACCACTTCACAAGTCGGAGGC |
| | | Reverse | CTGCAAGTGCATCATCGTTGTTC |
| | *il-12* | Forward | TTTATGATGGCCCTGTGCCT |
| | | Reverse | CAGCTCATCAATAACTGCCAGC |
| | *b2m* | Forward | ATTCACCCCCACTGAGACTG |
| | | Reverse | TGCTATTTCTTTCTGCGTC |
| Viral | *n* | Forward | CACATTGGCACCCGCAATC |
| | | Reverse | GAGGAACGAGAAGAGGCTTG |
